# Portal bile acids and microbiota along the murine intestinal tract exhibit sex differences in physiology

**DOI:** 10.1101/2024.07.12.603325

**Authors:** François Reichardt, Lauren N Lucas, Lydia Okyere, Jim Choi, Daniel Amador-Noguez, Christopher A Gaulke, Sayeepriyadarshini Anakk

**Author notes:** Co-Corresponding Authors Electronic address and.

## Abstract

Microbes in the intestine transform bile acids during transit, altering their functional and signaling capacities before absorption into the portal vein. Sex differences in the gut microbiota have been noted, but their consequence on bile acid composition is unclear. Here, we investigated the composition and imputed functional potential of microbes in the small and large intestine together with portal and systemic bile acids. Female and male mice exhibit distinct microbial diversity throughout the length of the intestine leading to dimorphism in genes related to bile acid transformation. Subsequently, we found that the total portal bile acid concentrations were doubled in female mice compared to males. Conversely, oxo-bile acids that are rare in systemic circulation represented almost 30 percent of the portal pool in the male mice. Oxo- and deconjugated-bile acids were absent in germ-free mice consistent with microbe-mediated bile acid transformation. More importantly, gnotobiotic mice do not show sex differences in portal bile acids. Taken together, we demonstrate that sex differences in gut microbiota cause dimorphism in bile acid levels within the enterohepatic loop.

**Highlights:**

- Murine microbial diversity exhibits sex-specific patterns along the small intestinal tract and colon.
- Distinct gut microbial profiles confer differential abundance of secondary bile acid metabolism genes in male and female mice.
- Portal bile acid compositions mirror sex differences in microbe-mediated bile acid processing that are lost in germ free conditions.

## Introduction

Bile acids (BAs) are cholesterol metabolites synthesized in the liver to facilitate fat absorption in the small intestine. Besides their amphipathic role, these C24 and C27 molecules function as hormonal signals to control various cellular processes and are efficiently recycled back to the liver through enterohepatic recirculation (*1-3*). Intestinal microbiota transform primary BAs (synthesized in the liver) into secondary BAs. Thus microbially-mediated BA modifications are fundamental in regulating the hydrophobic-hydrophilic balance of luminal BAs that subsequently affects dietary fat solubilization and signaling potential as different BAs target distinct receptors (*4-7*). Although occurrence of microbe-mediated deconjugation and oxidation of BAs in the upper part of small intestine along with lipid absorption was reported six decades ago (*8-11*), until recently most BA studies have focused on the ileum or the colon. The small percentage (∼5%) of luminal BAs entering the colon (*12-15*) underscores the significance of small intestinal microbes and their potential for BA modifications since microbiota have been noted all along the gastrointestinal tract in rodents and recently in humans (*16-19*). In addition, unexpected discoveries in microbe-mediated-BA modifications have led to a resurgence in interest in this field (*20-26*).

But, the regional distribution and sex differences of bacterial bile acid interplay have been overlooked. Therefore, in this study, we investigated the biogeography of the mucosal microbiota across the small and large intestine under normal physiology using both male and female wild-type mice. In both sexes, we quantified microbial taxonomic diversity and imputed functional potential in the small intestine (duodenum, jejunum and ileum), large intestine (cecum and colon), and feces. To decipher whether microbiota-mediated BA modifications occur in a sex-specific manner, we profiled the bile acid repertoires in portal blood to capture enterohepatic recirculation as well as systemic blood. Analogous experiments in germ free animals demonstrated that sex differences in BA composition are, at least in part, mediated by gut microbiota. Taken together, our findings demonstrate robust sex-differences in BA composition due to sex-specific variations in microbial diversity and uncover that oxo-BAs comprise a significant portion of the enterohepatic not the systemic BA pool.

## Methods

### Animals

Conventionally raised (CR) female and male C57Bl/6 mice were obtained from Jackson laboratories and maintained in flow cages in a temperature-controlled (23°C) facility on a standard 12-hour light/dark cycle. Female and male germ-free (GF) mice were bred in Rodent Gnotobiotic Facility, UIUC. All experiments were performed between 9 am -12 pm on 3-5 months old mice fed a normal chow *ad lib*. Euthanasia was performed by cervical dislocation. National Institute of Health guidelines for use and care of laboratory animals were followed, and all experiments were approved by the Institutional Animal Care and Use Committee at the University of Illinois at Urbana-Champaign (Champaign, IL, USA).

### Microbial community analysis

Microbiota analyses were done in a first group of CR mice from both sexes. Fresh feces were sampled before euthanasia. After collection, the duodenum (first 5 cm of the small intestine), jejunum (middle), ileum (last 5 cm), and proximal colon were delicately dissected in sterile ice-cold PBS before intestinal mucosa scraping with a microscope slide. Cecum was also collected. All samples were immediately frozen and stored at -80°C until processing. Microbial DNA was isolated using the DNeasy® PowerSoil® Pro Kit, Qiagen (Venlo, Netherlands) with the addition of a 10m incubation at 65°C prior to bead beating to facilitate cell lysis (*27*). The V4 region of 16S rRNA gene was then amplified in triplicate using golay barcoded primers (515F-806R) (*28*). Pooled triplicates were visualized on a 1% agarose gel and quantified using the Qubit® HS kit (Life Technologies, Carlsbad, CA, USA). Amplicon libraries were then pooled, cleaned using the QIAquick® PCR Purification Kit according to according to the manufacturer’s instructions, and sequenced on an Illumina MiSeq (Illumina; Foster City, CA) by the DNA Services Lab at University of Illinois Urbana Champaign Roy J Carver Biotechnology Center. The resulting 8,541,271 million 300-bp paired-end sequences were input into DADA2 (v.1.2.0) for quality filtering (truncLen = c(250, 200), maxEE = c(2,2), truncQ = 2, rm.phix = TRUE, maxN=0), sequence variant calling, chimera filtering (method = “consensus”) and taxonomic assignment against the Silva reference database (v138) (*29, 30*). Resulting sequence variant profiles were rarefied to a depth of 5,000 reads per sample (vegan::rrarefy) and alpha- and beta-diversity calculated using vegan and custom scripts. PICRUSt2 was used to impute the functional potential for each sample (picrust2_pipeline.py with default parameters) (*31, 32*). Bile acids transforming genes used in this analysis were filtered to consider protein families that were present in more than 5% of the samples and predicted to be encoded by at least one ASV, whose abundance varied by sex and tissue.

### Bile acids

Portal and systemic blood were collected under isoflurane anesthesia in CR and GF mice from both sexes. After exposition of the abdominal cavity, portal blood was collected with a syringe (diameter 0.46mm) advanced ∼2 mm into the portal vein in the direction of the intestines. Systemic blood from the same individual was collected in the seconds after portal sampling by intracardiac puncture before mouse euthanasia. All samples were immediately centrifuged (8000g, 3min) and serum was snap frozen. Once thawed, sera were mixed with methanol (1/4 ratio) before centrifugation (10 minutes, 14000rpm, 4C). BA composition was determined in the supernatants using ultrahigh-pressure liquid chromatography-tandem mass spectrometry (uHPLC-MS/MS) and system consisting of a Thermo Scientific Vanquish uHPLC system coupled to a heated electrospray ionization (HESI; using negative polarity hybrid quadrupole high-resolution mass spectrometer (Q Exactive Orbitrap; Thermo Scientific), as previously described (*21*, Supp. Table 1). Settings for the ion source were: auxiliary gas flow rate of 10, sheath gas flow rate of 30, sweep gas flow rate of 1, 2.5 kV spray voltage, 320°C capillary temperature, 300°C heater temperature, and S-lens RF level of 50. Nitrogen was used as nebulizing gas by the ion trap source. Liquid chromatography (LC) separation was achieved using a Waters Acquity UPLC BEH C18 column with 1.7 μm particle size, 2.1 × 100 mm in length. Solvent A was water with 10 mM ammonium acetate adjusted to pH 6.0 with acetic acid. Solvent B was 100% methanol. The total run time was 31.5 min with the following gradient: a 0 to 24 min gradient from 30% solvent B (initial condition) to 100% solvent B; held 5 min at 100% solvent B; dropped to 30% solvent B for 2.5 min re-equilibration to initial condition. The flow rate was 200 μL/min throughout. Other LC parameters were as follows: autosampler temperature, 4°C; injection volume, 10 μL; column temperature 50°C. Bile acid quantitation was achieved using standard concentrations of each bile acid ranging from 0.03125 to 4 μM to generate eight-point external standard curves. The detection limit was below 0.01 μM for all bile acids. Standards were purchased from Avanti Polar Lipids and dissolved and stored in methanol at -80 °C.

### Statistics

Associations between microbial diversity, sampling site, and sex were quantified using generalized linear mixed effects models (glmmTMB v1.1.5) using animal ID as the random effect (vegan::diversity and vegan:: specnumber; alpha diversity), and permutational multivariate analysis of variance (PERMANOVA; vegan::adonis2; beta diversity) using R (v4.1.1) and vegan (v 2.6.4). To quantify associations between individual amplicon sequence variants (ASVs) and sex or tissue we built two negative binomial generalized linear mixed effects models (glmmTMB v1.1.5). The first model included only the random effect of animal ID (i.e., the null model) while the second included the parameters sex and tissue as well as the random effect of animal ID (taxa ∼ sex + tissue + (1|Animal ID)). An analysis of variance (stats::anova, stats v4.1.1) was calculated to determine if the inclusion of sex and tissue improved model fit compared to our null model. False discovery rate was controlled using p.adjust(method = “fdr”). Differences in KEGG orthologous protein families were detected using generalized linear mixed effects models for families associated with secondary bile acid metabolism. The significance of model parameters were tested using Wald χ2 tests car::Anova (v3.1.1) and pairwise comparisons of the means was calculated using emmeans::emmeans (v1.8.5).

Bile acid composition data are expressed as percentages of total bile acids. Statistical analyses were performed using Mann–Whitney tests with Prism software (version 6.O7, GraphPad Inc, La Jolla, CA, USA) as Pearson’s correlations. P ≤ 0.05 was considered statistically significant.

## Results

### Microbial diversity varies throughout the intestines and is sex-specific

Bacterial transformations alter the hydrophobicity index of BAs and thus impact their micellar capacity for fat solubilization. We examined the mucosal bacteria along the intestines. As expected, feces (glmm; z = 11.7, p < 2.0×10^−16^), and the large intestine, including cecum (z = 9.1, p < 2.0×10^−16^) and colon (z = 12.8, p < 2.0×10^−16^) displayed significantly more diverse microbial communities (*33, 34*) compared to the small intestine (Fig. 1A, Supp. Table 2). Male mice exhibited increased bacterial richness compared to females throughout the intestine from the duodenum to the colon (z = 2.719, P = 0.007) (Fig. 1A). Microbiome beta-diversity (a measure of similarity for microbial communities) also varied significantly based on the intestinal region (PERMANOVA; R^2^ = 0.38, p = 2.0×10^−4^) and sex (R^2^ = 0.07, p = 2.0×10^−4^) (Fig. 1B, Supp. Fig. 1A-F). Consistently, the most abundant genera were regionally distinct. For example, the proximal small intestine was highly colonized by facultative anaerobic taxa belonging to the genera *Bifidobacteria* and *Lactobacillus* (*18, 35, 36*), while the distal portions of the gut were heavily colonized with obligate anaerobes belonging to *Lachnospiraceae* and genus *Faecalibaculum* (Fig. 1C). Since intestinal length and transit time can impact microbial communities, we quantified these parameters between male and female animals and did not find any significant differences (Supp. Fig 2 A-B), negating their role in facilitating the sex-specific microbial diversity.

**Figure 1.**
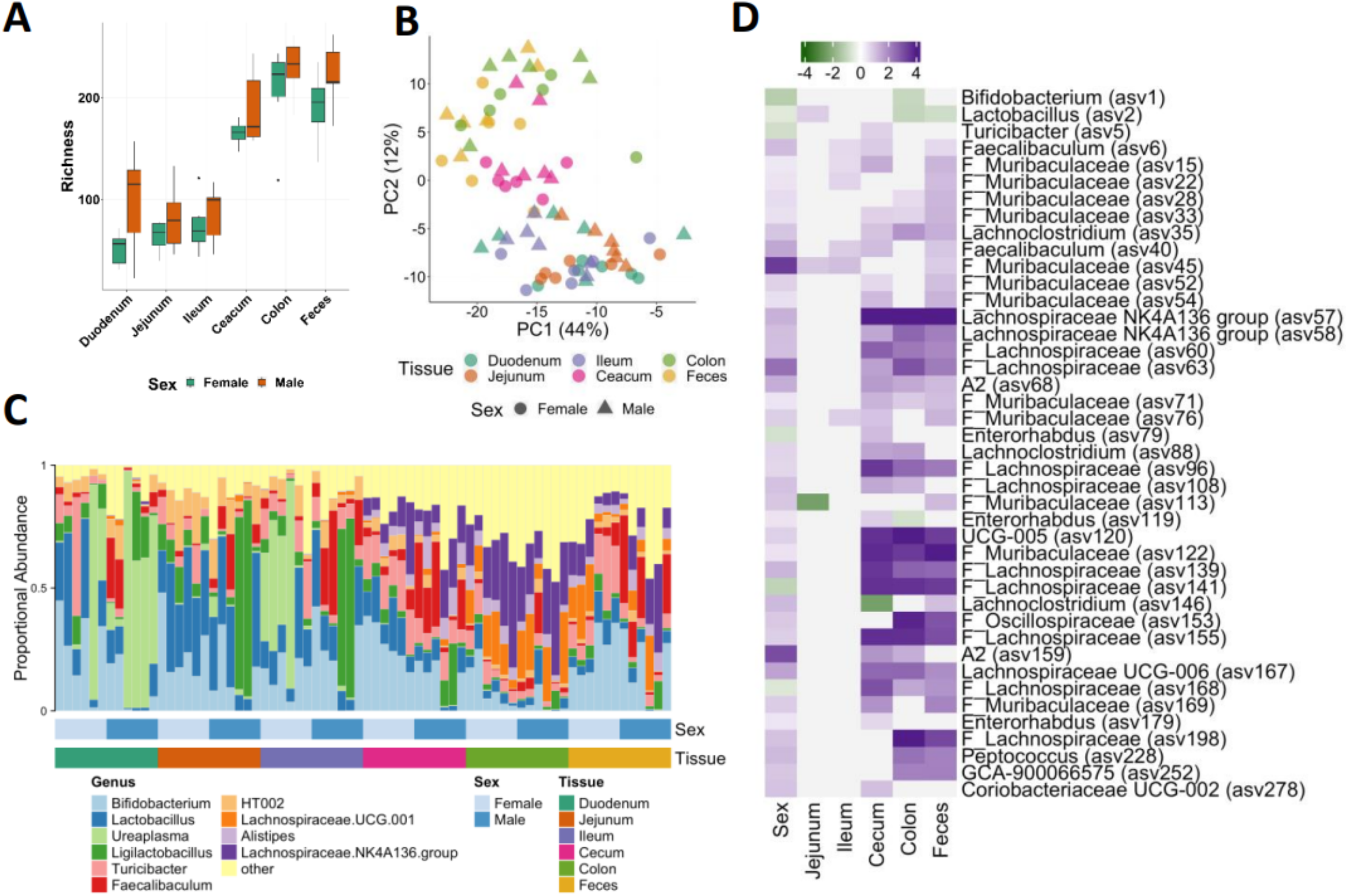
Abundance of bile acid producing bacteria vary by sex and gastrointestinal niche. A) Proportional abundance of the top 10 microbial genera across C57BL/6 gastrointestinal tissues. B) Microbial richness (ASV) in male and female mice along the length of the gut. C) Principal coordinate analysis of microbial beta-diversity (robust Aitchison distance) with sex indicated by point shape and tissue by point color. D) A heatmap of coefficients from generalized linear mixed effects models (q < 0.05) for which sex and tissue were observed to significantly impact amplicon sequence variant (ASV) abundance (p < 0.05). Color of each cell depicts the magnitude of the coefficient relative to female (sex) or duodenum (tissue). The genus level taxonomic assignment is provided when available, with the ASV identifier noted parenthetically, family level assignment (family prepended with F_) is provided when genus level assignment was not available. Grey cells indicate lack of significance (p > 0.05).

To precisely define the taxa that give rise to the sex- and region-based microbiome associations, we used robust negative binomial generalized linear mixed effects models. This approach identified 178 ASVs (FDR = 0.05) whose abundance was associated with either localization (118 ASVs) and or sex (60 ASVs). Most taxa that varied significantly between the sexes were increased in males relative to females including several important BA metabolizing groups (*e*.*g*., *Lachnospiraceae*). However, compared to females, males had reduced abundance of ASVs corresponding to the genera *Bifidobacterium* (asv1), *Lactobacillus* (asv2), *Turicibacter* (asv5), *Enterorhabdus* (asv79), and the family *Lachnospiraceae* (asv141 and asv168) (Fig. 1D). In line with our findings, Wu et al. noted higher levels of *Turibacter* and *Muribaculaceae* in small intestine of female mice. With respect to the location, we found a higher abundance of *Bifidobacterium* (asv1) and *Lactobacillus* (asv2) in the small intestinal compartment compared to the colon, while several ASVs corresponding to *Lachnospiraceae* and *Muribaculaceae* were increased in the large intestine and feces. Together these data demonstrate that both sex and location impact the abundance of microbial taxa capable of metabolizing bile acids.

### Sex differences in the intestinal microbiota converge on taxa that are BA metabolizers

Recent advances have demonstrated that the genes and pathways involved in bile salt metabolism are widely distributed across the bacterial tree of life (*21, 37, 38*). This means that large shifts in one taxa may not necessarily alter bile acid metabolism potential as long as the taxa being depleted are replaced with one or more that encode the same metabolic repertoire. To determine if the differential abundance of potential bile acid transformers noted above corresponded to differences in bile acid transformation machinery, we used PICRUSt2 to computationally impute the functional (gene) profiles of microbial communities between sexes and gut compartments. We constrained our analyses to protein families that are involved in bile acid metabolism and passed abundance and prevalence filtering (see methods; Kyoto Encyclopedia of Genes and Genomes; KEGG). After filtering we found three genes involved in bacterial BA metabolism: *bsh* (bile salt hydrolase) (K01442*), 7α-HSDH (hdhA*, K00076) and *baiN (*3-dehydro-bile acid delta 4,6-reductase, K07007). *Bsh* gene was encoded by a majority of taxa (64%), including *Bifidobacterium* and *Lactobacillus*, that are known to deconjugate BAs *in vivo* (*15*) but significantly varied by sex and location (Fig. 2A). More recently *bsh* is also shown to mediate acyltransferase activity and formation of amine-conjugated bile acids (*23, 24*). Overall, we find that BA transformation potentials found across the intestine and BA oxidation (*7α-Hsdh*, 15% of taxa) and 7α-dehydroxylation (baiN, 44% of taxa) are frequently encoded along with *bsh*, revealing diverse BA modification occurs more commonly than previously recognized. Moreover, the imputed gene abundances of *bsh* (χ2 = 203.9, p < 2.0×10^−16^), *hdhA* (χ2 =64.1, p = 1.7×10^−12^), *baiN* (χ2 = 256.4, p < 2.0×10^−16^), and total BA-associated genes (χ2 =32.4, p = 4.8×10^−6^) - the sum of *bsh, hdhA*, and *baiN* relative abundance - are significantly distinct within the intestinal niche (Fig. 2B-E). For instance, the highest relative *bsh* abundance was noted in the small intestine highlighting the relevance of this regions in BA metabolism. In line with our data, recent studies reveal comparable BSH expression between stomach, jejunum and cecum of the mice (*17*) and in fact, analysis of human samples also recapitulated the higher small intestinal *bsh* abundance (*18*).

**Figure 2.**
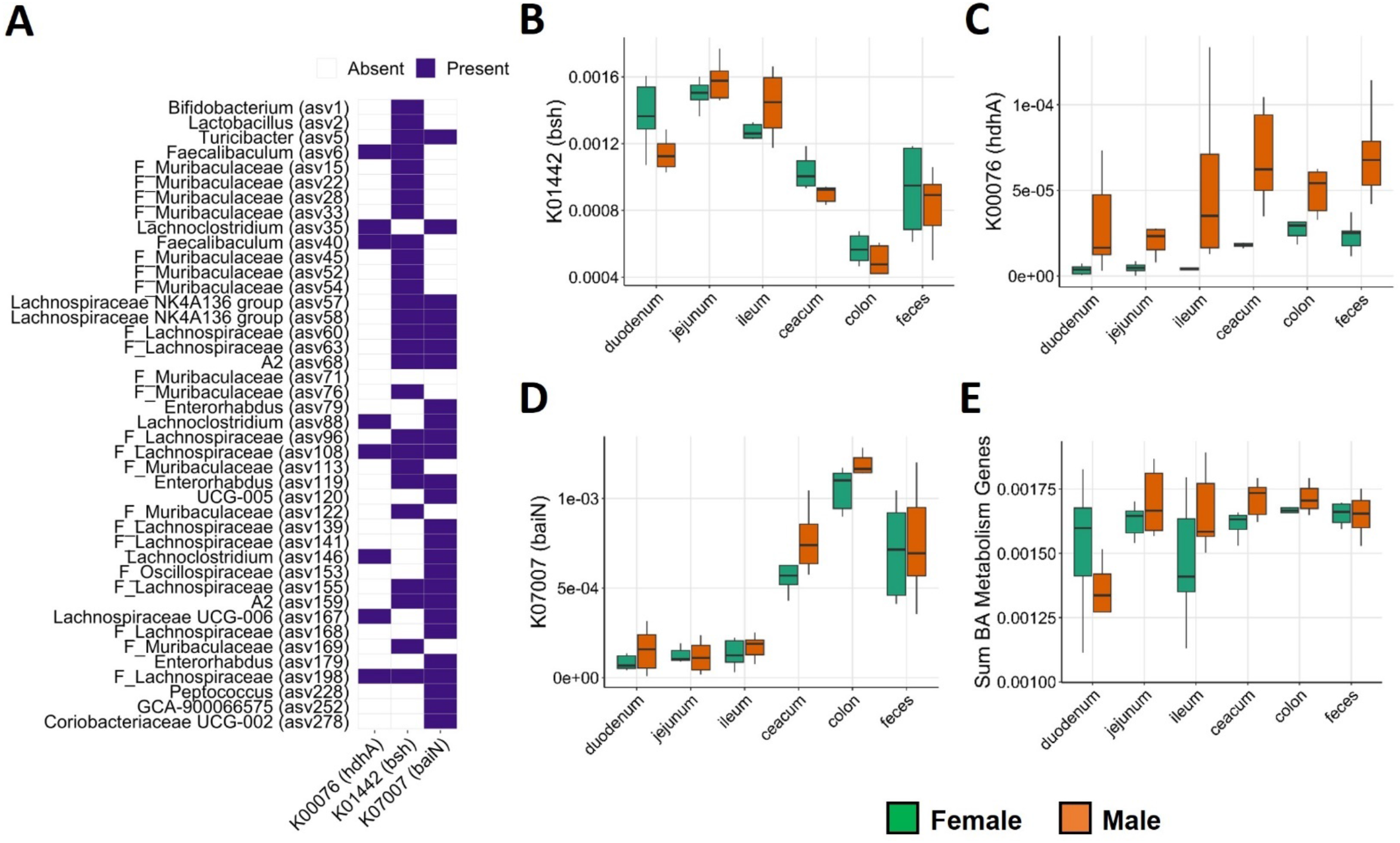
Bile acid modification potential elevated in male mice. A) Predicted presence of select bile acid metabolism gene families in ASVs which varied by sex and location (q < 0.05). Purple color indicates that the KEGG orthologous protein gene family is predicted to be encoded in the genome of this ASV, white indicates its predicted absence. The genus level taxonomic assignment is provided when available, with the ASV identifier noted parenthetically, family level assignment (family prepended with F_) is provided when genus level assignment was not available. The predicted relative abundance of B) bsh, C) hdhA, and D) baiN orthologs in imputed metagenomes in females (green) and males (orange) and E) the summed abundance of all genes associated with the secondary bile acid metabolism KEGG pathway (ko00121).

Distal portions of the small intestine exhibited increased abundance of *baiN, hdhA*, and total BA genes (Fig. 2C-E). Consistently, BA oxidation by 7α-HSDH was reported in human and rodent intestine highlighting its pervasive expression profile (*39-41*). We uncovered significant sex difference in the abundance of murine *hdh*A gene with higher levels in males versus females (χ2 = 44.5, P = 2.5×10^−11^) (Fig. 2C). Moreover, the summed abundance of *bsh, baiN* and *hdhA* genes showed sex-specific association (χ2 = 15.5, P = 8.1×10-3) such that the secondary BAs metabolism was higher in male mice (from jejunum to feces) irrespective of the niche (Fig. 2E).

Until now, due to overall microbial richness, microbe-BA-modifications have been primarily examined in the colon or feces. In contrast, our findings and other recent studies (*18, 42*) uncover that microbiota that mediate BA-transformations are present in the small intestines and are not confined to the colon or cecum. Given that BA reabsorption and feedback to the liver predominantly occurs in the ileum, it is not surprising that BA transformations may occur well before the large intestine. Investigating the enterohepatic recirculation (via the portal vein) will reveal several of the microbial-mediated BA modifications that occur.

### Portal BA composition differed between the sexes

Portal sampling captures BAs that are efficiently recycled back (∼95%) to the liver from the intestines, while systemic capture the excess that is spilled from the liver. By cannulating the portal vein, we obtain a snapshot of bacterial-BA modification that occurs in the intestine. We measured BA profiles from portal and systemic circulation from both sexes of WT mice (Supp. Table 3). We focused on primary BAs [βMCA, CA, CDCA] and microbe-dependent secondary BAs [UDCA, ωMCA, oxoBAs, deOH-BAs] in their free- or taurine-conjugated forms. Although not significant, total BAs in systemic circulation tended to be higher in females (P = 0.058) (Fig. 3A). Similar increases have been reported in female mice compared to males (*43*). In contrast, systemic BA composition was comparable between females and males except for 12-oxoLCA was higher in females, while LCA was higher in males (Fig. 3C-D).

**Figure 3.**
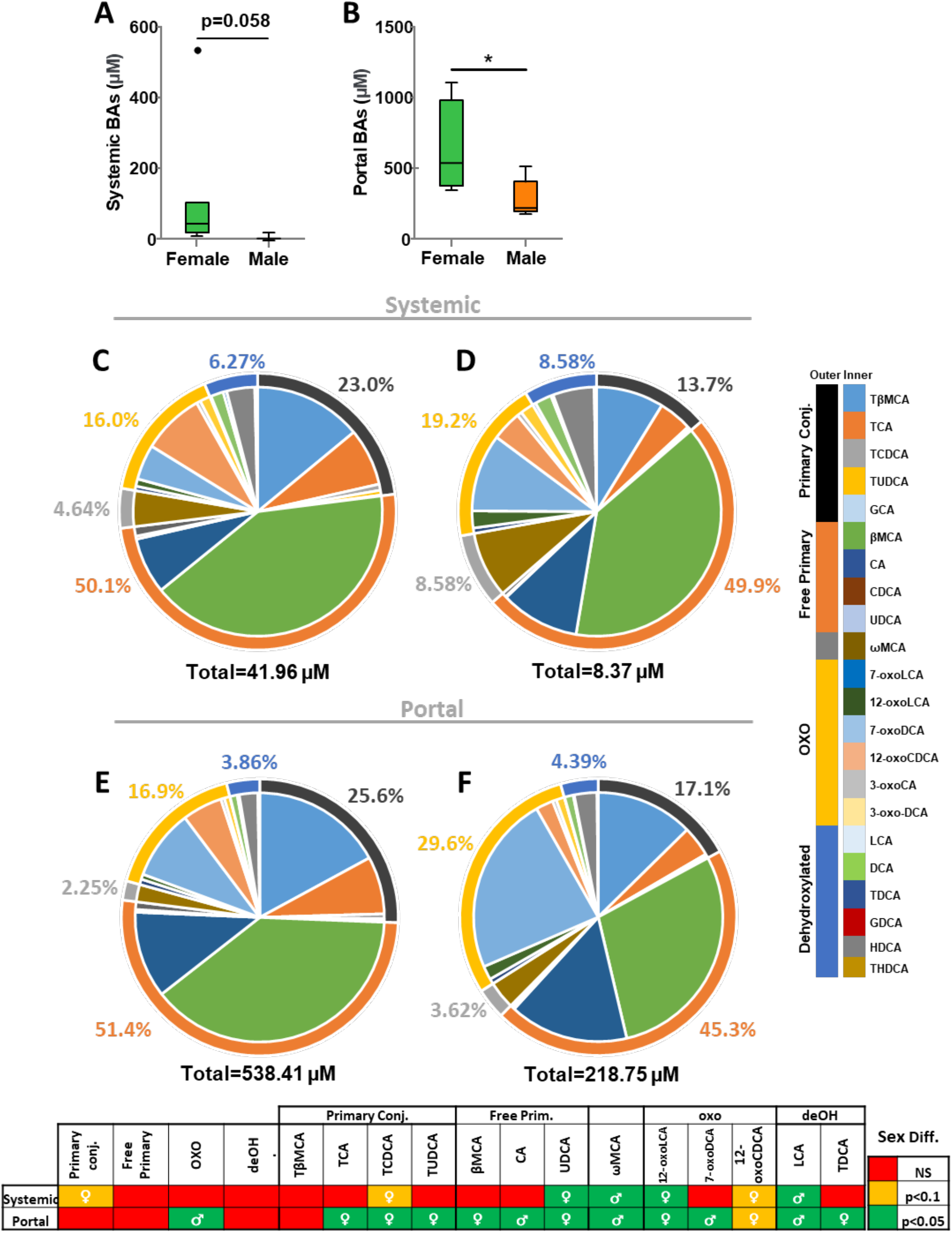
Increased bile acids oxidation in CR male portal but not systemic circulations. A) Systemic and B) portal BAs concentrations are elevated in females compared to males. Although females (C) and males (D) exhibit similar systemic compositions, portal compositions differ with increased oxo bile acids and 7-oxoDCA in males (F) compared to females (E). Portal: N=4♀/5♂; systemic: N=7♀/3♂.

As expected, portal BA concentrations were significantly higher than systemic levels (fold change ∼13X for females and ∼25X for males) (Fig. 3B). Moreover, female mice exhibited double the portal BAs and showed high amounts of conjugated BAs (TCA, TCDCA and TUDCA) (Fig. 3E-F). Free primary BAs and de-OH BAs were identical between the two sexes, except for higher UDCA levels were observed in female mice (Fig. 3E-F). In contrast, portal oxoBA levels are almost doubled in males, accounting for approximately 30% of total portal BAs, and only 16% in females. Of note, the three-fold increase in 7-oxoDCA in portal BA in males compared to females (23.3% vs. 7.9%), correlated with higher *hdhA* abundance we found in males (Fig. 3E-F).

Intriguingly, we uncover that male livers are exposed to high levels of 7-oxoDCA via the portal circulation on a regular basis in normal physiology, albeit its high systemic levels being reported only in liver, and gastrointestinal disease states (gall stones, hepatocellular carcinoma, MASLD, and IBD) (*37, 44-51*). Moreover, we do not find 7-oxoDCA in healthy murine colon content reflective of an efficient reabsorption via the portal vein (*41, 52*). We conjecture this occurs because the liver is adept at reducing 7-oxoDCA to CA (*53*), one of the major primary BA and thus deters its systemic spilling into the circulation.

Unlike males with smaller 12-oxoCDCA levels, female portal circulation displayed comparable proportions of 7-oxoDCA and 12-oxoCDCA (Fig. 3E-F). This result clearly demonstrates sex-specific oxidation of BAs. Additional experiments will be needed to determine why male mice exhibit a preference for oxidation of BAs at carbon 7 vs.12, whereas oxoBAs at both carbon positions are equally seen in females. Nevertheless, portal 7-oxoDCA and 12-oxoCDCA positively correlated with CA but inversely with TCA (Supp. Fig. 3A-F). 12-oxoCDCA is used as a precursor for UDCA synthesis (*54*), and in fact we find this correlation as well (Supp. Fig. 3G). Recent data (26, 57), also suggest oxo-BAs may locally act to alleviate intestinal damages in a sex-specific manner.

Nonetheless, there are limitations with studying intestinal luminal BAs as it does not account for passive diffusion/transport, and similarly hepatic and systemic BAs are snap shots and do not differentiate between enterohepatic recycling, *de novo* synthesis and transport. For examples, we find portal TCA is 3.9% whereas in the liver it is 35-40% in male mice (*55*). Thus, portal measurements mirror microbial BA biotransformation, and account for the transport across the intestinal mucosa without the hepatic *de novo* synthesis.

### Gut microbiota direct the sex differences in portal BA composition

We finally evaluated if the observed differences in portal BAs stem from gut microbiota-mediated BA transformation. Adult GF mice from both sexes exhibit longer small intestines and colon compared to CR animals (Supp. Fig 2A). Experiments were performed in an identical manner in the GF mice, wherein we collected systemic and portal samples for BA analysis. As expected, unconjugated primary BAs and secondary BAs were negligible in the serum and the portal circulation of GF mice (Figure 4C-F). Strikingly, the amount of total BAs in portal circulation in gnotobiotic female mice dropped to half of what was noted in CR female mice, bringing it down to the same level observed in GF male mice. Total portal BA levels were unchanged between CR and gnotobiotic male mice. Both sexes showed reduced concentrations of total systemic BAs as previously reported (*3, 56*). Unlike CR mice that have bMCA as a predominant BA, the gnotobiotic animals have T-bMCA as a major proportion both in systemic and portal circulation irrespective of the sex (Fig. 3E-F, 4E-F). Although absolute concentration of systemic and portal BAs in female GF mice were lowered, their differences were similar (∼13X), highlighting that BAs spill from liver into systemic circulation irrespective of composition. Also, we find that male GF mice show more unconjugated bMCA than their female counterparts. Although it is a rather small proportion (∼3% in systemic and 0.5% in portal), it implies that some deconjugation is feasible without microbes in the GF males. Importantly, there were no sex differences between the gnotobiotic mice that is consistent with oxo-BAs being generated by the bacterial BA transformations. These data clearly demonstrate that the sex differences in BA composition are driven by the gut microbiota.

**Figure 4.**
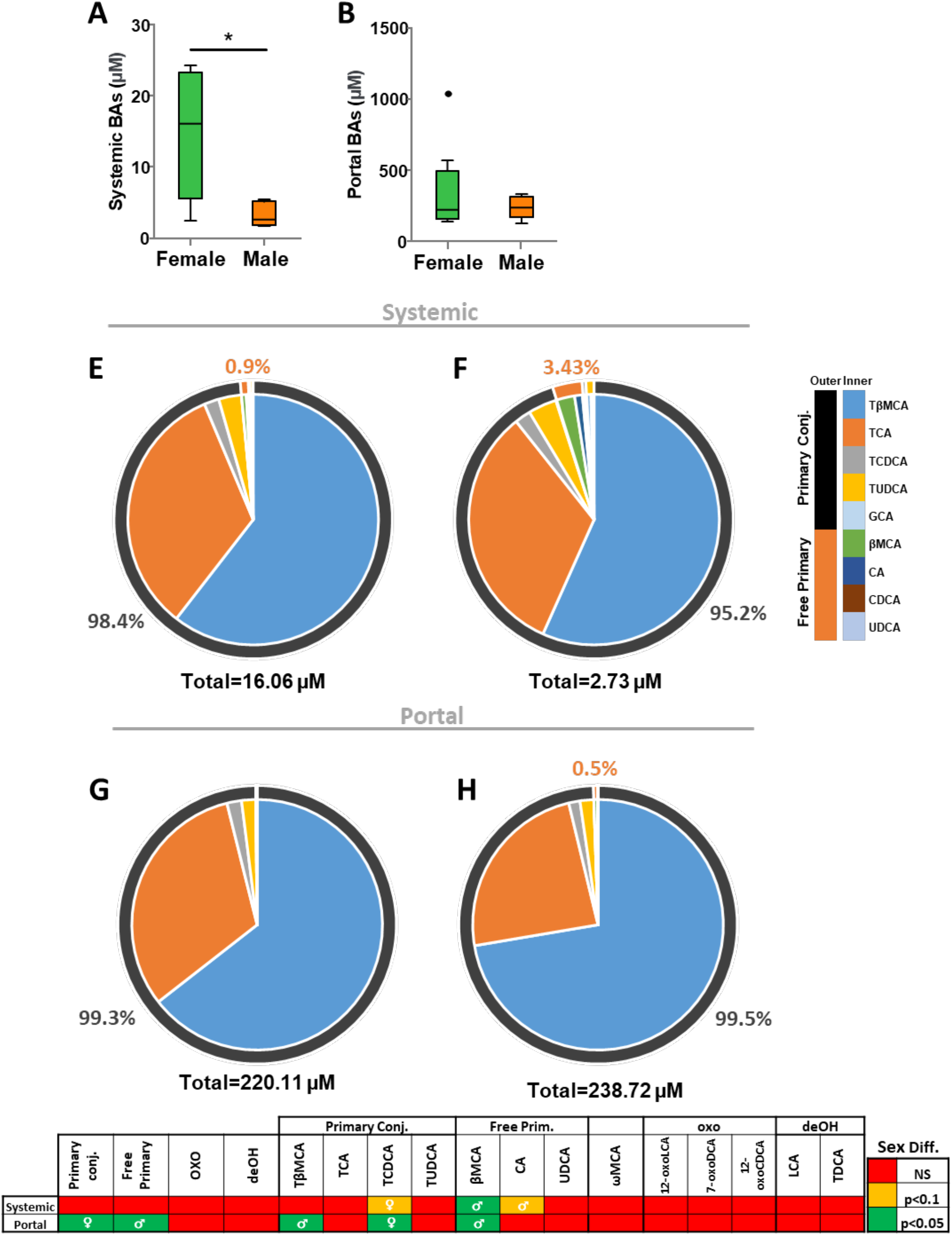
Microbiota is essential for sex differences in oxo-BAs. A) Systemic but not B) portal BAs concentrations are elevated in females compared to males. Females and males exhibit similar systemic (C and D) and portal (E and F) compositions, with no **s**econdary bile acids.

## Conclusion

Our findings uncover that microbiota across the entire gut, from duodenum to the colon, exhibit varied ability to transform BAs. Not only are these BA modifiers niche-specific but also sex-specific. Importantly, this is the first study that tapped into the portal vein to assess enterohepatic recirculation immediately after BA transformation in both sexes of adult mice. BA-microbiota interactions are bidirectional, so by utilizing GF mice we provide strong evidence that loss of microbiota is sufficient to abolish sex differences in portal BA composition and concentration in CR mice. We speculate that diet, age, and hormones would directly influence the microbe-BAs continuum. It is intriguing that the dramatic sex differences seen in portal BAs are not found at the systemic level. These findings imply that although female and male livers may receive different BA species back from the gut, these differences are nullified such that hepatic BA pools look comparable. In line with this idea, several prior studies evaluating hepatic BAs, including ours, do not find obvious sex differences (*55, 58*). We also speculate that such portal BA diversity, along with growth hormone and sex steroid hormones, may contribute to establish the inherent sex differences in liver metabolism and function. Finally, the sex differences in microbiota across the gastrointestinal tract may help explain certain diseases of the intestines and the liver that exhibit gender disparity. Nevertheless, we do not exclude the possibility that our mucosal samples may contain luminal bacteria which could also transform BAs.

## Abbreviations

3-oxoCA: 3-oxocholic caid
3-oxoDCA: 3-oxodeoxycholic acid
7-oxoDCA: 7-oxodeoxycholic acid (= 7-ketodeoxycholic acid)
7-oxoLCA: 7-oxolithocholic acid
12-oxoCDCA: 12-oxochenodeoxycholic acid (= 12-ketodeoxycholic acid)
12-oxoLCA: 12-oxolithocholic acid
BA: bile acid
bai: bile acid inducible operon
βMCA: β-muricholic acid
CA: cholic acid
CDCA: chenodeoxycholic acid
DCA: deoxycholic acid
FXR: Farnesoid-X-Receptor
GCA: glycocholic acid
HDCA: hyodeoxycholic acid
HSDH: hydroxysteroid dehydrogenase
LCA: lithocholic acid
PICRUSt: Phylogenetic Investigation of Communities by Reconstruction of Unobserved States
ω-MCA: ω-muricholic acid
TβMCA: tauro-β-muricholic acid
TCA: taurocholic acid
TCDCA: taurochenodeoxycholic acid
THDCA: taurohyodeoxycholic acid
TUDCA: tauroursodexycholic acid
UDCA: ursodeoxycholic acid.

## Acknowledgement

American Cancer Society, Nationale Institute of Health, Cancer Center of Illinois

## Authors contributions

FR, LNL, LO, JC were all responsible for the acquisition of the data, DN, CG, and SA were involved in the design and execution and analysis along with FR, LNL, LO, and JC. All authors provided critical feedback and helped shape the research, analysis, and manuscript.

