## Supplementary figures for "Portal bile acids and microbiota along the murine intestinal tract exhibit sex differences in physiology"

**Supplemental Material**

*GI transit* 4-months-old CR mice were overnight fasted before to be gavaged with 0.1ml of a methylene blue solution (50mg methylene blue / 1ml of PBS + 1% carboxy methyl cellulose). Dye progression was measured in the small intestine 1h after gavaging.

**Supplemental Figures**

**Supp. Figure 1. Sex differences in microbial diversity all along the intestines of CR mice.** Principal coordinate analysis of microbial beta-diversity (robust Aitchison distance) with sex indicated by point color in A) duodenum, B) jejunum, C) ileum, D) ceacum, E) colon and F) feces. Duodenum, ceacum and feces: P<0.05. Jejunum, ileum and proximal colon: P<0.1.

**Supp. Figure 2. Intestinal length is longer in GF mice from both sexes compared to CR.** Compared to CR mice, both GF females and males exhibit longer small intestine (A). CR mice from both sexes exhibit analogous upper GI transit (B). Transit: N=5♀/6♂.

**Supp. Figure 3. Variations in portal BA compositions.** A) Portal oxo-BAs correlate with Primary Conj. BAs. B-C) Correlations between TCA, CA and 7-oxoDCA suggesting that the increase of 7-oxoDCA in males depend on the availability of CA following TCA deconjugation. E-F) Correlations between TCA, CA and 12-oxoCDCA. G) Positive correlation between 12-oxoCDCA and UDCA. H). Correlation almost significant between portal 7-oxoDCA and 12-oxoCDCA. CR mice, N=5♀/5♂.

**Supplemental Tables**

**Supp. Table 1. Bile acid names and structure description for standards used for LC-MS/MS analysis.**

**Supp. Table 2. Microbiota composition (contrast_table). (Attached as Excel Sheet)**

**Supp. Table 3. Bile acid compositions. (Attached as Excel Sheet)**

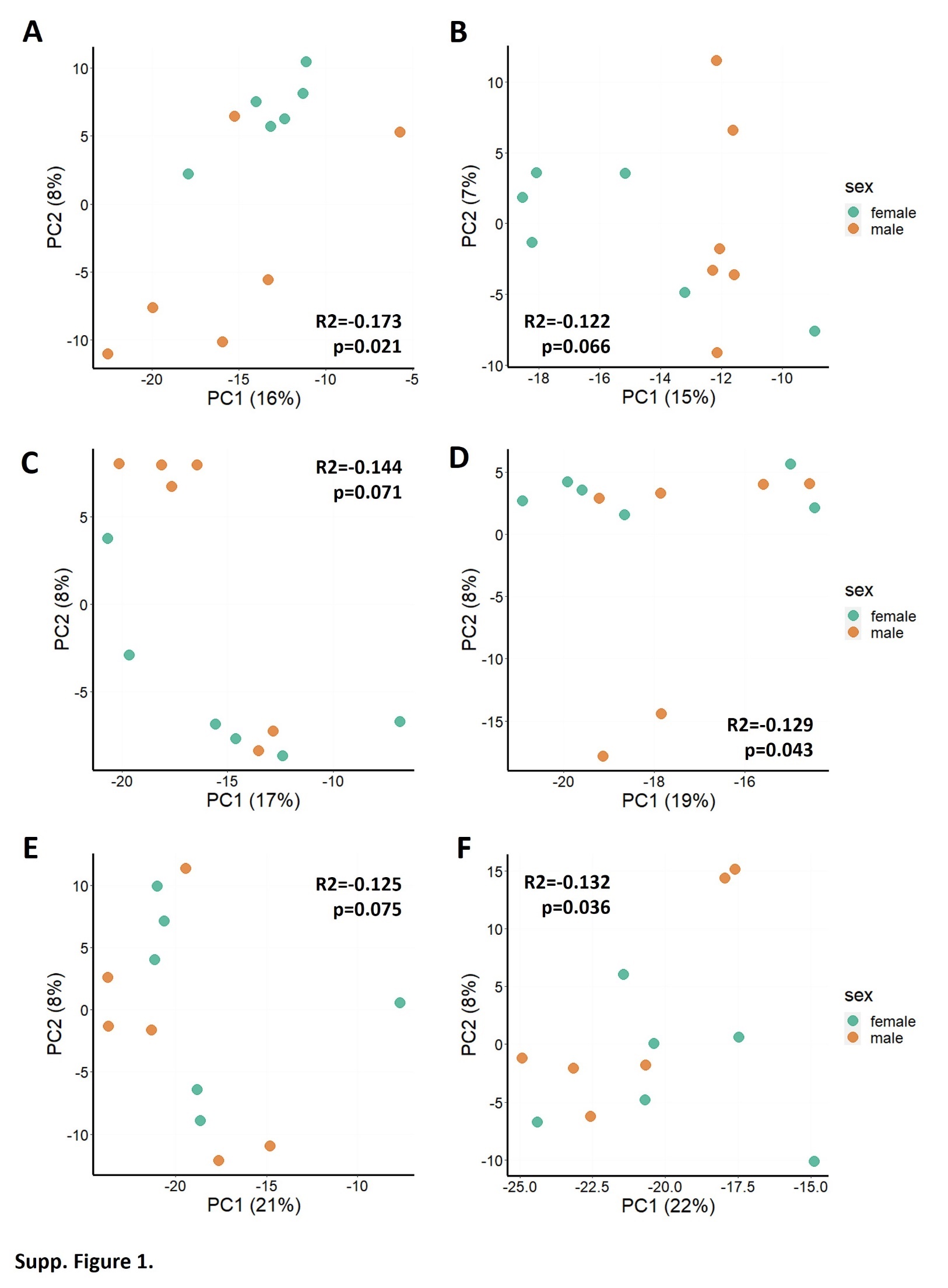

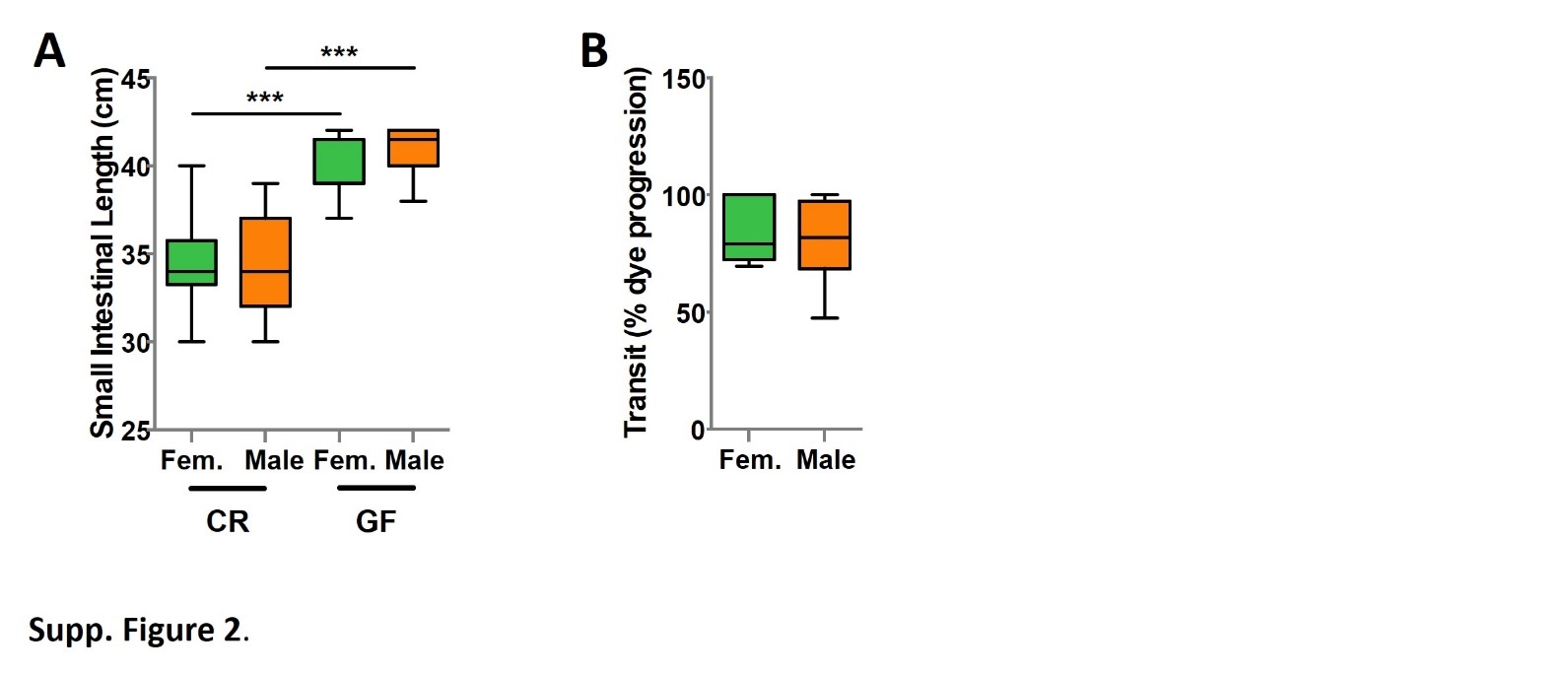

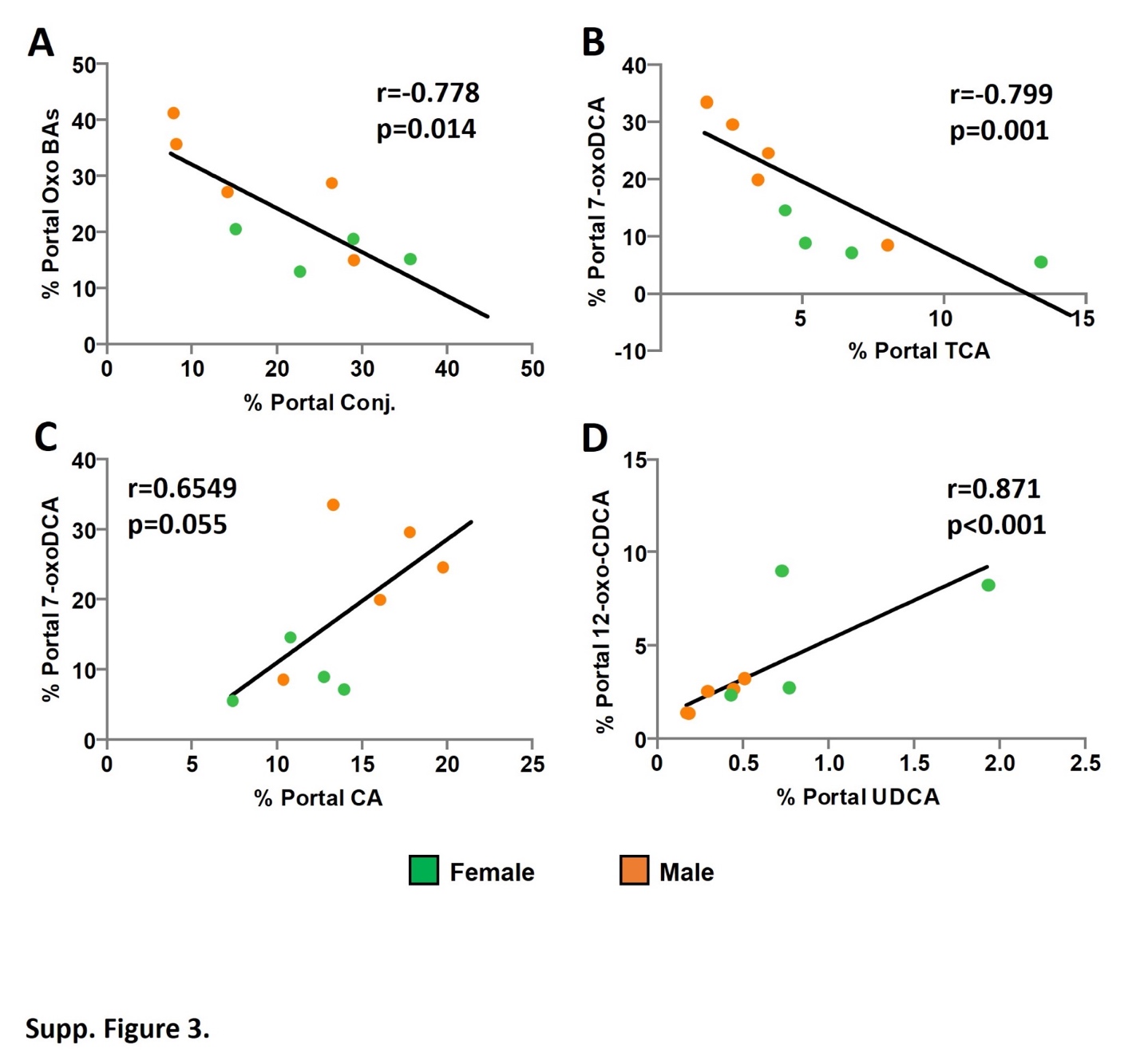

| **Supplemental Table 1. Bile acid names and structure description for standards used for LC-MS/MS analysis.** | | | | | | | | | | | |
| --- | --- | --- | --- | --- | --- | --- | --- | --- | --- | --- | --- |
| **Bile Acid Common Name** | **Abbreviation** | **Mol. Formula** | **C1** | **C2** | **C3** | **C6** | **C7** | **C12** | **DB** | **R** | **Rt** |
| 1.      Glycocholic acid | GCA | C26H43NO6 | α,β-H | α,β-H | α-OH | α,β-H | α-OH | α-OH | N/A | G | 14.05 |
| 2.      Taurocholic acid | TCA | C26H45NO7S | α,β-H | α,β-H | α-OH | α,β-H | α-OH | α-OH | N/A | T | 13.9 |
| 3.      Cholic acid | CA | C24H40O5 | α,β-H | α,β-H | α-OH | α,β-H | α-OH | α-OH | N/A | H | 15.04 |
| 4.      Glycochenodeoxycholic acid | GCDCA | C26H43NO5 | α,β-H | α,β-H | α-OH | α,β-H | α-OH | α,β-H | N/A | G | 15.94 |
| 5.      Taurochenodeoxycholic acid | TCDCA | C26H45NO6S | α,β-H | α,β-H | α-OH | α,β-H | α-OH | α,β-H | N/A | T | 15.73 |
| 6.      Chenodeoxycholic acid | CDCA | C24H40O4 | α,β-H | α,β-H | α-OH | α,β-H | α-OH | α,β-H | N/A | H | 17.1 |
| 7.      Glycoursodeoxycholic acid | GUDCA | C26H43NO5 | α,β-H | α,β-H | α-OH | α,β-H | β-OH | α,β-H | N/A | G | 12.51 |
| 8.      Tauroursodeoxycholic acid | TUDCA | C26H45NO6S | α,β-H | α,β-H | α-OH | α,β-H | β-OH | α,β-H | N/A | T | 12.45 |
| 9.      Ursodeoxycholic acid | UDCA | C24H40O4 | α,β-H | α,β-H | α-OH | α,β-H | β-OH | α,β-H | N/A | H | 13.75 |
| 10.   Glycodeoxycholic acid | GDCA | C26H43NO5 | α,β-H | α,β-H | α-OH | α,β-H | α,β-H | α-OH | N/A | G | 16.32 |
| 11.   Taurodeoxycholic acid | TDCA | C26H45NO6S | α,β-H | α,β-H | α-OH | α,β-H | α,β-H | α-OH | N/A | T | 16.17 |
| 12.   Deoxycholic acid | DCA | C24H40O4 | α,β-H | α,β-H | α-OH | α,β-H | α,β-H | α-OH | N/A | H | 17.46 |
| 13.   Glycolithocholic acid | GLCA | C26H43NO4 | α,β-H | α,β-H | α-OH | α,β-H | α,β-H | α,β-H | N/A | G | 17.75 |
| 14.   Taurolithocholic acid | TLCA | C26H45NO5S | α,β-H | α,β-H | α-OH | α,β-H | α,β-H | α,β-H | N/A | T | 17.61 |
| 15.   Lithocholic acid | LCA | C24H40O3 | α,β-H | α,β-H | α-OH | α,β-H | α,β-H | α,β-H | N/A | H | 19.09 |
| 16.   Hyocholic acid/γ-Muricholic acid | HCA/γ-MCA | C24H40O5 | α,β-H | α,β-H | α-OH | α-OH | α-OH | α,β-H | N/A | H | 13.7 |
| 17.   Tauro-α-muricholic acid | T-α-MCA | C26H45NO7S | α,β-H | α,β-H | α-OH | β-OH | α-OH | α,β-H | N/A | T | 10.51 |
| 18.   α-muricholic acid | α-MCA | C24H40O5 | α,β-H | α,β-H | α-OH | β-OH | α-OH | α,β-H | N/A | H | 11.83 |
| 19.   Tauro-β-muricholic acid | T-β-MCA | C26H45NO7S | α,β-H | α,β-H | α-OH | β-OH | β-OH | α,β-H | N/A | T | 10.65 |
| 20.   β-muricholic acid | β-MCA | C24H40O5 | α,β-H | α,β-H | α-OH | β-OH | β-OH | α,β-H | N/A | H | 12.03 |
| 21.   ω-muricholic acid | ω-MCA | C24H40O5 | α,β-H | α,β-H | α-OH | α-OH | β-OH | α,β-H | N/A | H | 11.89 |
| 22.   Glycohyodeoxycholic acid | GHDCA | C26H43NO5 | α,β-H | α,β-H | α-OH | α-OH | α,β-H | α,β-H | N/A | G | 12.77 |
| 23.   Taurohyodeoxycholic acid | THDCA | C26H45NO6S | α,β-H | α,β-H | α-OH | α-OH | α,β-H | α,β-H | N/A | T | 12.94 |
| 24.   Hyodeoxycholic acid | HDCA | C24H40O4 | α,β-H | α,β-H | α-OH | α-OH | α,β-H | α,β-H | N/A | H | 14.34 |
| 25.   3-oxocholic acid | 3-oxoCA | C24H38O5 | α,β-H | α,β-H | =O | α,β-H | α-OH | α-OH | N/A | H | 13.41 |
| 26.   7-oxodeoxycholic acid | 7-oxoDCA | C24H38O5 | α,β-H | α,β-H | α-OH | α,β-H | =O | α-OH | N/A | H | 11.83 |
| 27.   12-oxochenodeoxycholic acid | 12-oxoCDCA | C24H38O5 | α,β-H | α,β-H | α-OH | α,β-H | α-OH | =O | N/A | H | 12.15 |
| 28.   3-oxochenodeoxycholic acid | 3-oxoCDCA | C24H38O4 | α,β-H | α,β-H | =O | α,β-H | α-OH | α,β-H | N/A | H | 15.83 |
| 29.   3-oxodeoxycholic acid | 3-oxoDCA | C24H38O4 | α,β-H | α,β-H | =O | α,β-H | α,β-H | α-OH | N/A | H | 16.13 |
| 30.   Isolithocholic acid | isoLCA | C24H40O3 | α,β-H | α,β-H | β-OH | α,β-H | α,β-H | α,β-H | N/A | H | 17.27 |
| 31.   7-oxolithocholic acid | 7-oxoLCA | C24H38O4 | α,β-H | α,β-H | α-OH | α,β-H | =O | α,β-H | N/A | H | 14.13 |
| 32.   12-oxolithocholic acid | 12-oxoLCA | C24H38O4 | α,β-H | α,β-H | α-OH | α,β-H | α,β-H | =O | N/A | H | 14.63 |
| 33.   Taurodehydrocholic acid | T-dhCA | C26H39NO7S | α,β-H | α,β-H | =O | α,β-H | =O | =O | N/A | T | 5.98 |
| 34.   Glycodehydrocholic acid | G-dhCA | C26H37NO6 | α,β-H | α,β-H | =O | α,β-H | =O | =O | N/A | G | 5.87 |
| 35.   Isodeoxycholic acid | isoDCA | C24H40O4 | α,β-H | α,β-H | α,β-H | α,β-H | α-OH | α-OH | N/A | H | 18.49 |
| 36.   6,7-dioxolithocholic acid | 6,7-dioxoLCA | C24H36O5 | α,β-H | α,β-H | α-OH | =O | =O | α,β-H | N/A | H | 14.08 |
| 37.   7,12-dioxolithocholic acid | 7,12-dioxoLCA | C24H36O5 | α,β-H | α,β-H | α-OH | α,β-H | =O | =O | N/A | H | 8.18 |
| Rt = retention time. R represents side chain abbreviations: G = glycine, T = taurine, H = hydrogen. DB = double bond. | | | | | | | | | | | |
| For numbered carbons on steroid core, see Figure 2. | | | | | | | | | | | |
